# Comparison of B values obtained by two methodologies that use soil as a substrate and their influence on the estimation of the proportion of nitrogen fixed by legumes

**DOI:** 10.1101/2022.06.17.496592

**Authors:** Verónica Berriel, Jorge Monza, Carlos Perdomo

## Abstract

The B value is required to quantify the nitrogen derived from the atmosphere (%Ndfa) in the Rhizobium-legume symbiosis using the ^15^N natural abundance method. When the B value of a particular specie is not known, one possibility Is to use as a proxy the B value of a specie from the same genus, but this can cause the estimate of %Ndfa to be inaccurate. In this work, we compared two methodologies for determining the B value of *Crotalaria juncea, C. spectabilis, C. ochroleuca* and *Cajanus cajan*, using soil as the substrate. One method involved growing plants in soil and averaging the lowest δ^15^N values of plant shoots (B-minimum), while the other consisted in adding sucrose to soil to immobilize the mineral nitrogen (N-immobilized), and then averaging the shoot δ^15^N values of all plants. Results showed that B values of *C. cajan* and *C. ochroleuca* obtained using the N-immobilized method were up to 1‰ lower than those reported in the literature for these species. Therefore, we propose that, at least in these species, B values determined with the N-immobilized method should be used to estimate the%Ndfa.

## Introduction

Cover crops (CCs) based on species of the genus *Crotalaria* and *Cajanus cajan* can contribute with significant amounts of nitrogen (N) to the soil through the biological fixation process (BNF) (Giller, 2001; Santana et al., 2018, Berriel et al. 2020). The high BNF of these CCs is achieved due to the efficient symbiotic association that they establish with a wide variety of rhizobia soil strains of the genus *Methylobacterium, Bradyrizobium* and others (Sy et al., 2001; Zilli et al., 2020, Fossou et al., 2020).

The accurate quantification of the N fixed by these CCs is needed to balance N inputs, and this information constitutes an important input for planning sustainable agricultural rotations (Landriscini et al., 2019).

Estimation of the N proportion in plants or crops that derives from the atmosphere by BNF (%Ndfa) can be carried out using the nuclear technique proposed by Shearer and Kohl (1986), which is based on the natural differences in ^15^N that exist between the atmosphere and the soil (Amarger et al., 1979). The determination of%Ndfa by this technique is carried out via Equation 1 (Unkovich et al., 2008).

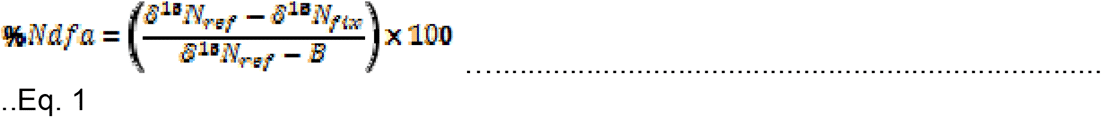

where δ^15^N_ref_ is the ^15^N abundance determined in a non-fixing reference plant, δ^15^N_fix_ is the ^15^N abundance of the legume that has grown and developed in the site and/or soil of interest, and B is the ^15^N abundance of the legume whose only N source is the atmospheric N.

A key component in Eq. [1] is the B value, which in some cases could be obtained from the literature. However, it should be noted that this value depends on the plant species (Unkovich et al., 2008) and the rhizobium strain with which symbioses were established (Guimarães et al., 2008; Pauferro et al., 2010).

Usually, the B value is determined by growing plants inoculated with a rhizobium strain on inert substrates (sand, vermiculite, hydroponic solutions, etc.), providing the necessary requirements for plant growth, except mineral N. Although the growth may be limited under these conditions, this ensures that the only N source available of for the plant is the atmospheric N (Unkovich et al., 2008).

Another strategy to estimate the B value is to perform a sampling of a site where the species of interest is growing and estimate as value B an average of the lowest. Here, the mean of the most negative δ^15^N shoot values is used as the Bestimate (B-minimum), assuming that in these cases the plant’s N came exclusively from the BNF (Peoples et al., 2002). A different approach to determine B is force the immobilization of mineral N from the soil (N-immobilized) by adding a substrate with a high C/N ratio, in such a way that the plant does not have N from the soil for its growth and that this mineral N does not inhibit BNF (Ferguson et al., 2019). A material highly energetic or with a high C/N ratio promotes net N immobilization (Mori et al., 2012; Romero et al., 2015), because soil microorganisms will require more N to metabolize the added C (Chen et al., 2020). Therefore, soil nitrate and ammonium will be converted into organic forms and incorporated into cellular components, such as proteins, thus leaving the soil devoid of mineral N (Cao et al., 2020).

Moreover,these two approaches can also be used with uninoculated seeds, which allows to estimate the B value of the rhizobia strains already present and adapted to this particular soil. Thus, this B value will be representative of a specific soil- rhizobia-legume symbioses system.

Our aim was to compare the B value obtained with these two methods that use soil as the substrate not only between themselves, but also with values reported in the literature for the same species, and to assess the impact of this variations in the%Ndfa estimations.

## Materials and Methods

### Plant material and growing conditions

Plants of *Crotalaria juncea, Crotalaria spectabilis, Crotolaria ochroleuca* and *Cajanus cajan* were cultivated in pots containing samples of an agricultural soil (4kg pot^-1^) which has never been planted with these legume species. Before planting, seeds *were* superficially sterilized (Okito et al., 2004), and sown at a rate of one seed per pot. The soil was an Argiudol from southern Uruguay (organic carbon = 11.6 g/kg; sand = 24.5; silt = 48.7%; clay = 26.8%; NO_3_ ^-^: 3.6 mg/kg; NH_4_ ^+^:_3 4_7.1 mg/kg).

Plants were grown for 90 days in a growth chamber at 30ºC, with a variable relative humidity of between 30 and 50% and a light intensity of 500 µmol m^-2^. s^-1^ with a 16/8 h light/dark cycle and irrigated with deionized water.

To determine the B value according to the minimum B method, plants of each specie were grown in 40 pots containing the original soil.

To determinate the B value with the N-immobilized method, soil was mixed with sucrose (in ratio 1 kg:5 g) and then incubated at 30ºC for 20 days, after which the soil mixture was divided among sixteen pots, four pots for each plant species.

### Biomass production and analytical measurements

After harvesting, samples of leaves, stems, roots, and nodules from each plant were dried separately at 60 °C until constant weight and then the dry mass of each part was determined. All samples were first ground with a fixed and mobile blade mill (Marconi MA-580) until reaching a particle size of less than 2 mm, and then with a rotary mill (SampleTek 200 vial Rotator) until reaching the required granulometric size for isotopic analysis. Samples were weighed into tin capsules, and their total N concentration and ^15^N natural abundance were then determined in a Thermo Finnigan DELTAplus mass spectrometer (Bremen, Germany) coupled to a Flash EA 1112 elemental analyzer through a ConFloIII interface.

The isotopic ratio was expressed in delta notation (δ) in parts per thousand (‰) using Equation 2 (Sulzman, 2007):

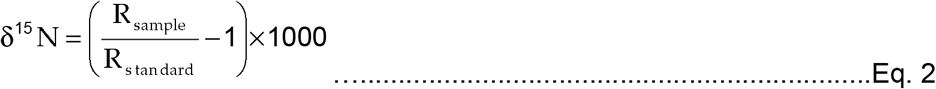

where R is the ratio of intensities (measured in the mass spectrometer) of the least abundant to the most abundant isotope.

Equations 3, 4 and 5, respectively, were used to determine the δ^15^N values for the shoot, radicular system, and the whole plant of each replication:

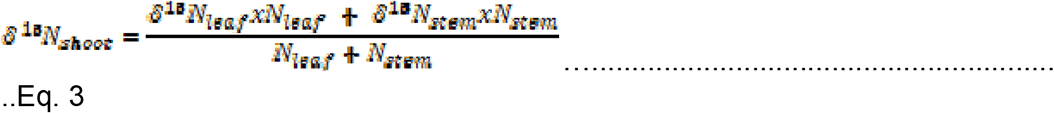

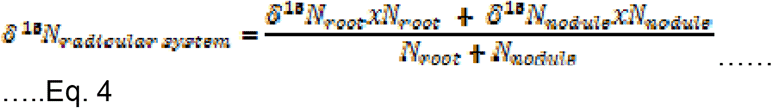

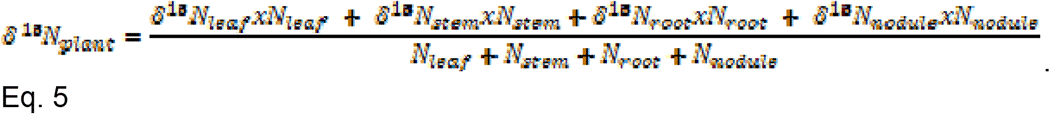

where N-leaf, N-steam, N- and N-nodule are the amounts in grams of N in the leaf, stem, root and nodule, respectively.

The δ^15^N shoot value (B value) for the minimum method was obtained as the average of the four lowest δ^15^N shoot values, while that for the N-immobilized method as the average of all δ^15^N values.

To estimate the %Ndfa plants from each specie were grown in six additional pots under the same conditions as in the B-minimum method assay. The mean δ^15^N value from these 6 pots was used as the value of δ^15^N-fix in Equation 1. These estimations were made not only with the two B values determined in this work, but also with those reported in the literature. Reference plants was maize with δ^15^N = 9.7 **‰**.

### Experimental design and statistical analysis

We studied the effect of two factors (B value determination method and species) on the N mass and ^15^N isotopic composition of each plant part (leaf, stem, root, nodules) and their partial totals (part area, root system) or the total plant, with a completely randomized statistical design.

The species factor had four levels (*C. juncea*, C. *spectabilis, C. ochroleuca* and *C. cajan*) and the determination method had 2 levels (B-minimum, N-immobilized) The main effects (method and species), as well as Their interaction was analyzed by Anova, but when the interaction was significant, the effect of the method was compared by Anova only within each species. The means were compared by LSD, and the existence of significant differences was assumed when p≤ 5%.

## Results

### Total N mass in different plant parts

In all species, the highest N mass in the shoot was found in leaves (approximately 83%), while in the root system each of its components contributed differently depending on the species. In *C. spectabilis* and *C. ochroleuca* the contribution from the nodule and the root to the N mass entire root system was 40 and 60%, respectively; while in *C. juncea* these contributions were 60 and 40%, respectively. On the other hand, in *C. cajan* the nodule and the root contributed 20 and 80% of the total N mass of the root system, respectively.

For the N mass of leaves, no interaction was found between the main factors species x method, and significant differences were found between both species and methods (Fig. 1 A). *Cajanus cajan* accumulated more N than the *Crotalaria* species. Regarding the method, more N was accumulated when immobilized-N method was used, compared to minimum-B method, except in *C. ochroleuca*, where there were no differences neither for leaf nor for aerial part. In stems, on the other hand, the N mass was influenced by the interaction between the species and the methods, for which the method effect was analyzed separately for each species (Fig. 1B). As expected, in all species the total aerial part followed the same trend as the leaf, given the greater contribution of the N mass in leaves to this total Fig. 1 C).

**Fig. 1.**
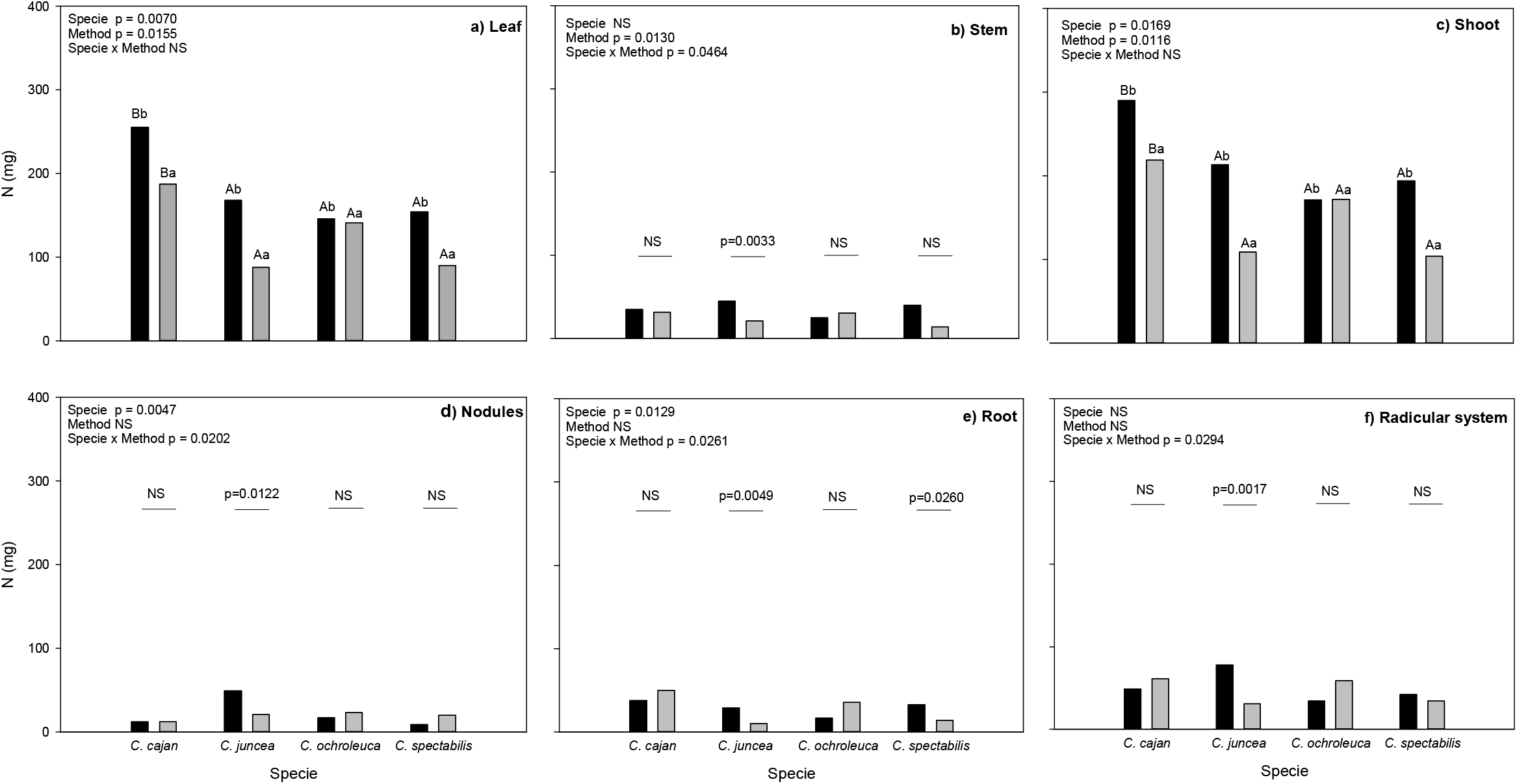
Masa de N en diferentes partes de las plantas que fueron cultivadas sobre distinto sustrato o métodos para determinar el valor B (N-inmovilizado y B-mínimo). Las barras negras y grises indican la masa de N obtenida con el método de N-inmovilizado y B-mínimo respectivamente. Las letras mayúsculas y minúsculas indican las diferencias entre especies y métodos respectivamente (p<0,05).

As in the aerial part, the N mass of nodules and roots was influenced by the species-by-method interaction; thus, the method effect was analyzed separately for each species. In nodules, only in *C. juncea* the N mass was different between methods in (Fig. 1 D), while in roots there were differences in both *C. juncea* and*C. spectabilis* (Fig. 1 E), but the total root system followed a trend that was more similar to that observed in nodules (Fig. 1 F). As occurred for leaves and shoot, when there were significant differences between methods, in all species the highest N mass accumulation was registered with the immobilized-N method.

For the root system, the species with similar trends were grouped together; one group was formed with *C. cajan* and *C. ochroleuca*, and the other with *C. juncea* and *C. spectabilis*, but in none of these groups the method-by-species interaction was significant. In addition, within both groups, the results were maintained, since the method was only significant in the first group and the higher values were obtained with the immobilized-N method.

In the whole plant, the N mass followed a similar same trend as in the leaf, because this part had contributed with most N to the shoot, and in turn, the N in shoot constituted on average 78% of the total. As it had happened in leaf, the species and method effects were statistically significant, but the species-by-method interaction was not. The mass of N accumulated in the whole plant was higher in*C. cajan* compared to the *Crotalaria* species, and again, the highest N mass was obtained with the minimum-B method (Table 1).

### δ^15^N

For both the total of the aerial part and the root system, the δ^15^N value was estimated from the weighted addition of its parts (leaf and stem on one side and nodules and root on the other), as was indicated in Eqs. 4 and 5. In all species, leaves had δ^15^N values that were closer to zero (less negative) than in stems, and since leaves contained more N mass, shoot δ^15^N values were similar to those of leaves. The δ^15^N values of the root system, on the other hand, were always positive and between those of nodules and roots, since the N mass of both parts was similar and their δ^15^N values were also positive, although more positive in nodules than in roots.

In the case of the δ^15^N values of leaves and stem, the shoot components, a significant species-by-method interaction was found (Fig. 2A and 2B, respectively). Thus, separate ANOVAS for each species revealed that only in leaves of *C. juncea* and *C. cajan* did the δ^15^N values vary between methods (Fig. 2A). In stems, on the other hand, a similar result was observed in *C. juncea* and *C. spectabilis* (Fig. 2B). In both cases, when the results were statistically significant, the most negative values were always recorded for the immobilized-N method.

**Fig. 2.**
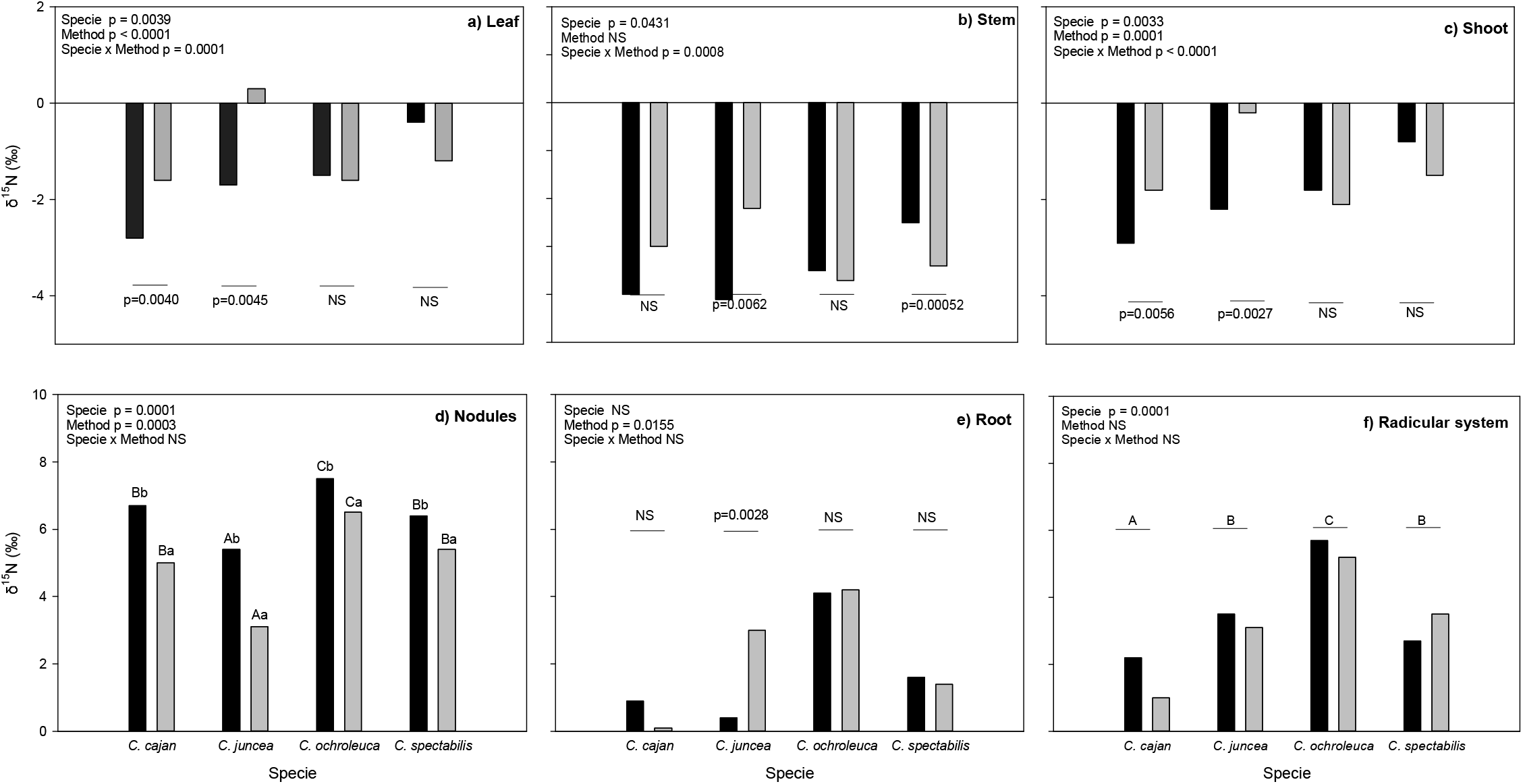
Valores de δ15N en diferentes partes de las plantas que fueron cultivadas sobre distinto sustrato o métodos para determinar l valor B (N-inmovilizado y B-mínimo). Las barras negras y grises indican la masa de N obtenida con el método de N-inmovilizado y B-mínimo respectivamente. Las letras mayúsculas y minúsculas indican las diferencias entre especies y métodos respectivamente (p<0,05).

The δ^15^N values of shoots followed the same trend as its constituent parts, resulting in a significant method-by-species interaction of the ANOVA (Fig. 2C). This analysis is, however, of special importance, because the δ^15^N value of the shoot also represents the B value of the plant. The results of the separate ANOVAs for each species showed in *C. cajan* and *C. juncea* that B values differed between methods, being also more negative when the immobilized-N method was used. In the other two species (*C. ochroleuca* and *C. spectabilis*), instead, the method did not significantly affect the B value.

Due to this interaction between method and species, the B value differences between species were evaluated only within each method. For the minimum-B method, significant differences (p=0.0033) were detected; and mean comparisons showed that *C. juncea* (-0.18 ‰) differed from the rest of the other species, which in turn did not differ among themselves (-2.06; -1.77 and -1.46 ‰, for *C. ochroleuca, C. cajan* and *C. spectabilis*, respectively). On the other hand, for the immobilized-N method, there were also significant differences between the Bvalues of the species (p<.0001). Mean comparisons showed that B values of *C. juncea* and *C. ochroleuca* were statistically equal (-2.21 and -1.83‰, respectively), but they were different from those of *C. cajan* (-2.94‰).) and *C. spectabilis* (0.94‰), which in turn also were different from each other.

In nodule and root, there was no significant interaction between method and species (Fig. 2D and 2E). In nodules, statistical differences were found both between both species and methods (Fig. 2D). In root, the ^15^N isotopic composition was different between methods only in *C. juncea*, with less positive values in the immobilized-N method (Fig. 2E). The root system showed a different tendency to its parts, since differences were only found at the species level (Fig. 2F). This result was obviously the consequence of the mass balance between the sum of the parts of nodules and roots.

In the whole plant, the ^15^N isotopic composition of all species followed a similar trend as that of the leaf, due to the greater contribution of the N mass of this part to the total plant. There was no significant interaction between species and method, but the main effects of species and method were significant. In terms of species,the δ^15^N values were more negative in *C. cajan* than in the other species, while, in terms of method, the δ^15^N values were more negative with immobilized-N (Table 1).

### Proportion of N derived from atmosphere

Estimations of %Ndfa obtained with Eq. 3 by substituting with the B values obtained with the minimum-B or immobilized-N methods, revealed that the greatest difference were obtained in *C. juncea* (13%) and *C. cajan* (8%); in these cases, the lower %Ndfa values were obtained with immobilized-N method. In addition, %Ndfa values obtained with previously published B values were also higher than those obtained with minimum-B in *C. cajan, C. juncea* and *C. ochroleuca* by 14; 8.6 and 8.5%, respectively (Table 2).

## Discussion

### N mass derived from atmosphere and growth media

The two methods for B value determination produced different levels of N mass derived from fixation because the BNF process was enhanced when plants grew on soil with sucrose addition. This result was expected, since the incorporation to the soil of plant residues with a high C/N ratio, such as corn or rice straw, has already been used to enhance soil mineral N immobilization and increase BNF (Mori et al., 2012; Salgado et al., 2021), and sucrose is an energetic material with no N in its chemical structure.

The inverse relationship between the Ndfa mass in the evaluated legumes and the mineral N availability in the substrate was consistent with the fact that these species are native to tropical areas with limited N availability and legumes evolved to depend on their BNF potential to acquire N (Trytsman et al., 2019; Jaiswal and Dakora, 2019). Thus, these species are particularly suitable to be used as CC in environments with low N availability (dos Santos Nascimento et al., 2021; Chu et al., 2004; Fan et al., 2006). To this respect, *C. cajan* was in this study the species with the highest BNF capacity, even without specific rhizobia inoculation, depending only in the native soil strains.

The distribution pattern of δ^15^N among the different plant parts was similar in all the evaluated species, being the shoot depleted while the root enriched in ^15^N. This pattern was consistent with reports from other species that grew on substrates without mineral N inputs (Gathumbi et al., 2002; Okito et al., 2004; Woldekirstos et al., 2014). In *Crotalarias*, instead, the δ^15^N composition of the entire plant was close to zero, indicating that although isotopic fractionation occurred within plant parts, it did not happen during the BNF process itself (Unkovich, 2013). Conversely, the ^15^N isotopic composition of *C. cajan* at the whole plant level was negative, a result that had also been reported for other tropical legume species (Okito et al. 2004; Unkovich, 2008; Woldekirstos et al., 2014).

With respect to these cases, Unkovich (2013) interpreted that they could be the result of error accumulations, spanning from culture conditions to analytical processing. Chalk and Croswell (2018), in turn, disagreed with at least part of this interpretation, since the δ^15^N variability of interlaboratory rounds is generally low. Although these last authors did not elaborate any further, it should be noted that if these negative δ^15^N values were just the result of randomness, positive values should also have been reported, but instead they fail to show up in the literature. At least in our study, the whole-plant negative δ^15^N values found in *C. cajan* but not in *Crotolarias* would not have been the result of different growing conditions or analytical processing, since all evaluated species were grown and processed similarly. Therefore, it would be possible that there were real ^15^N isotopic fractionation differences during BNF between these two groups.

### Influence of B value determination methods

The minimum-B methodology was originally proposed to estimate the %Ndfa in white clover and ryegrass pastures grazed by cattle at open-air, receiving, thus, animal depositions (Hansen and Vinther, 2001). In these situations, the δ^15^N values of the mixed pasture (grasses and legumes) were more negative than the B values determined in sand-vermiculite media, resulting in the estimation of negative %Ndfa values. This reduction of the isotopic values was due to the absorption by plants of N from the urine, which was δ^15^N depleted with respect to that of the original mixed pasture This depletion occurred during animal metabolism, with δ^15^N variations that ranged from -1.7‰ in the original pasture to -2.8 ‰ in the urine (Steele & Daniel, 1978).

Some of this deposited N is then volatilized as NH_3_, which is further depleted in ^15^N. The absorption of this N by leaves would explain the δ^15^N decreases of grasses from positive values to -7‰, cited by Eriksen and Høgh-Jensen (1998). On the other hand, the nitrogenous compounds remaining in the soil become ^15^N enriched (Robinson, 2001). To this regard, Tonn et al. (2019) applied urine (δ^15^N=2‰) to a mixed pasture of *L. perenne* and *T. repens*, and found that *L. perenne* leaves were initially rapidly depleted in ^15^N compared to a control without this application (δ^15^N 0.1 vs. 5.8‰, respectively). The leaves of *T. repens*, on the other hand, did not change their isotopic composition. Subsequently, foliar δ^15^N values increased (*T. repens* to 4.5‰ and *L. perenne* to 5.9‰), presumably due to the absorption of these remaining inorganic soil compounds enriched in ^15^N. At the end of the experiment, δ^15^N changes were higher in grasses than in legumes, in part because the ammonia-N absorption rate by leaves tend to be inversely proportional to its foliar N concentration, which is lower in grasses (Tonn et al., 2019)). In addition, this result could be partly a consequence of the BNF process, which tends to imprint in legumes the δ^15^N value of air.

Based on the cited results from other authors, as well as those found in this study, we consider that prior to making %Ndfa estimates, it would be advisable to compare the B values to be used with the ^15^N signature of the plant material, especially for legumes under grazing. Mori et al. (2012), evaluated different methodologies to determine B values for *Lotus corniculatus, Trifolium repens*, and *Trifolium pretense*, including, among others, the immobilized-N and minimum-B methods. These authors observed that the B values obtained with the immobilized- N method were always more negative than those obtained with the minimum-B method, although they were not statistically different from each other. In other words, this reported trend was similar to that found in our study for *C. cajan* and *C. juncea*, although in the case of *C. spectabilis* and *C. ochroleuca*, the B values identified from these two methods were similar (Fig. 2C).

To estimate the %Ndfa in *C. spectabilis* and *C. ochroleuca*, some authors have used the B values reported for other legumes of the same genus, because there were no published B values for these two species (Resende et al., 2002; Ojiem et al., 2007). In this work, we have determined specific B values for the symbiotic association between these two species and native soil rhizobia. In the case of *C. juncea* and *C. cajan*, on the contrary, several B values had already been proposed (Boddey et al., 2000; Gathumbi et al., 2002). In spite of that, our results suggest that even for these species it would be preferable to use specific B values locally estimated. This is because as suggested by Chalk et al. (2017) and Woldekirstos et al. (2014), B values would tend to vary with the native or inoculated Rhizobium strain and with the legume genotype or cultivar.

When comparing the %Ndfa calculated with the B values estimated with the immobilized-N method obtained in this work with those from the bibliography, differences were only relevant in *C. cajan* and *C. ochroleuca*, the species with the highest %Ndfa values. In other words, the importance of using specific B values increases as plants acquire a greater proportion of their N from the atmosphere. This explains why no practical %Ndfa differences were found in *C. juncea* when B values estimated with these two methodologies were used. Overall, we would recommend the immobilized-N method for determining local B values, since this methodology constitutes a simple and practical way to integrate into the B value the local effects of growing conditions and rhizobium strains.

---

**Cuadro 1.**
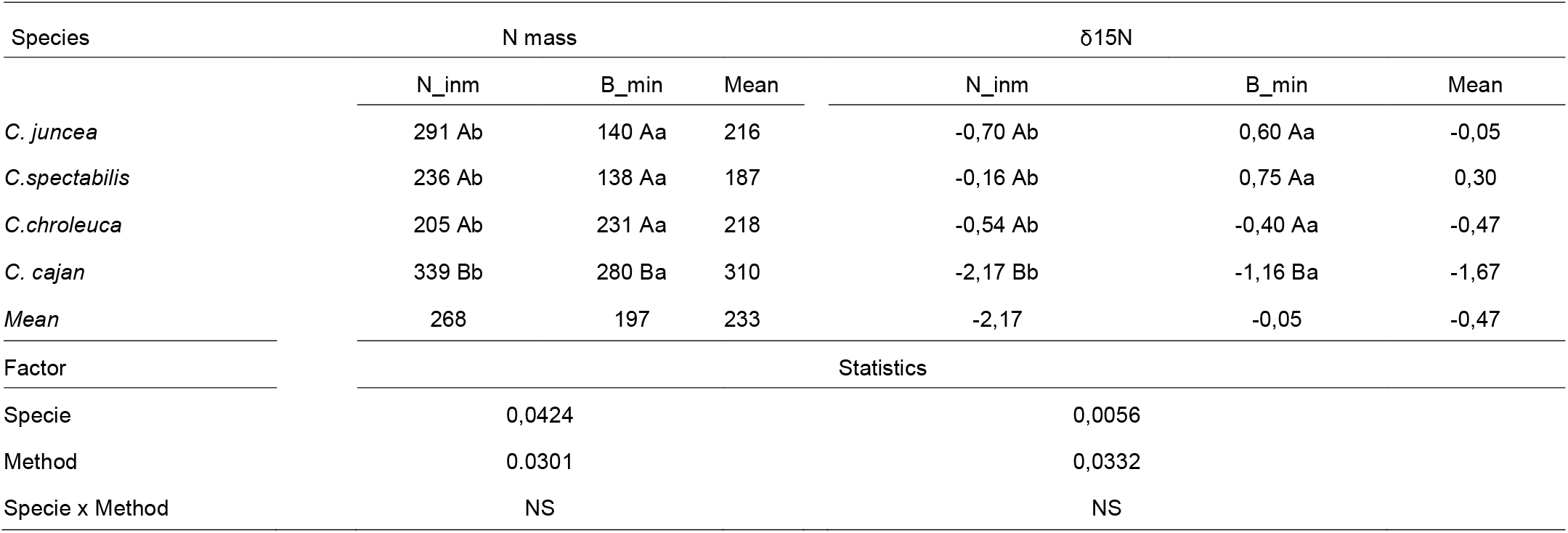
Valores medios de la masa de N (expresada en mg) y de δ15N (expresada en ‰) en la planta entera en cuatro especies de leguminosas obtenidos con dos métodos N- inmovilizado (N-inm) and B-mínimo (B-min). Las letras mayúsculas y minúsculas indican las diferencias entre especies y métodos respectivamente (p<0,05).

**Cuadro 2.**
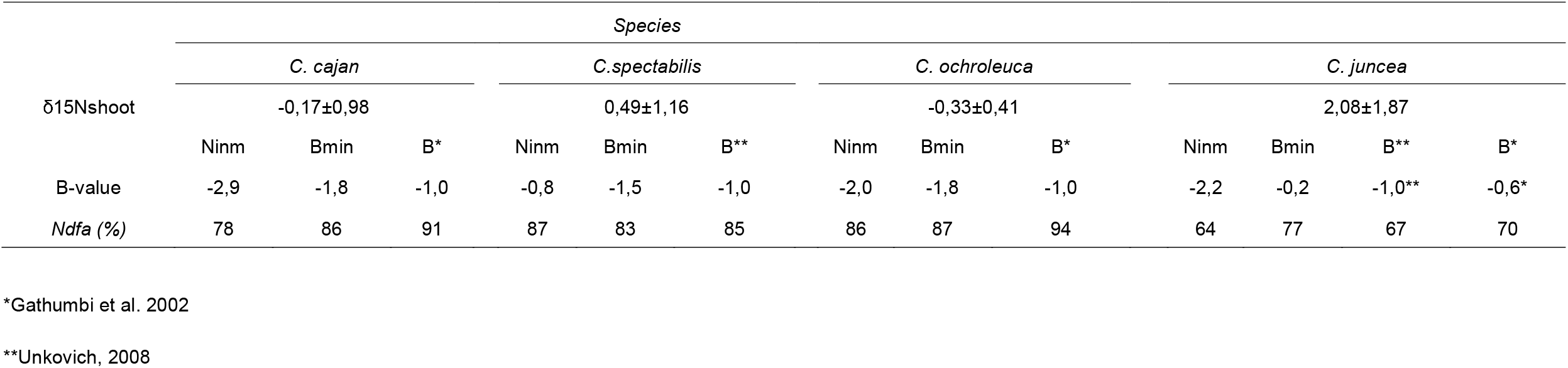
Proporción de nitrógeno fijado del aire (%Ndfa) usando valores B determinados en este estudio y otros de la literatura.

